# 17β-Estradiol Counteracts Pathological Microtubule Remodeling To Enhance Cardiac Function

**DOI:** 10.1101/2025.01.22.634271

**Authors:** Ryan Moon, Neal T. Vogel, Jenna B. Mendelson, Lynn M. Hartweck, John P. Carney, Minwoo Kim, Melissa K. Gardner, Sasha Prisco, Kurt W. Prins

## Abstract

The female-predominate sex hormone 17β-estradiol exerts cardioprotective effects via multiple mechanisms. Available data demonstrate 17β-estradiol modulates microtubule dynamics *in vitro*, but its effects on pathogenic microtubule remodeling in pressure-overloaded cardiomyocytes are unexplored. Here, we show 17β-estradiol directly blunts microtubule polymerization *in vitro*, counteracts endothelin-mediated microtubule remodeling in iPSC-cardiomyocytes, and mitigates microtubule stabilization in pulmonary artery banded right ventricular cardiomyocytes. 17β-estradiol treatment blunts cardiomyocyte and nuclear hypertrophy, restores t-tubule architecture, and prevents mislocalization of connexin-43 in RV cardiomyocytes of pulmonary artery banded rats. These cellular phenotypes are paired with significant improvements in RV function. Thus, we propose 17β-estradiol exerts cardioprotective effects via direct modulation of microtubules in addition to its well ascribed signaling functions.

---

Cardiomyocyte microtubule remodeling occurs in multiple species challenged by pressure overload, and heightened microtubule density promotes cardiomyocyte dysfunction^1^. The microtubule cytoskeleton impacts several areas of cardiac biology by regulating cardiomyocyte hypertrophy, nuclear remodeling, t-tubule structure, and the proper localization of the gap junction protein connexin-43^1^. In addition, cross-talk between the microtubule cytoskeleton and the desmin intermediate filament cytoskeleton regulates some of the microtubule-dependent cardiomyocyte phenotypes^1^. 17β-estradiol, the female-predominant hormone, induces microtubule depolymerization in breast cancer cells lacking estrogen receptors^2^. *In vitro*, 17β-estradiol slows microtubule polymerization rates^3^, which demonstrates 17β-estradiol directly regulates microtubule dynamics. While 17β-estradiol exerts cardioprotective effects via pleotropic mechanisms^4^, its effects on pathological microtubule remodeling in pressure-overloaded cardiomyocytes are unexplored. Here, we investigate the hypothesis that 17β-estradiol suppresses pathological microtubule remodeling, which restores cardiomyocyte/nuclear size, t-tubule morphology, and connexin-43 localization and ultimately enhances cardiac function in pulmonary artery banded rats.

A fluorescent tubulin kit (Cytoskeleton:BK011P) quantified *in vitro* microtubule polymerization rates in the presence of ethanol vehicle or 17β-estradiol. Human induced pluripotent stem cells (iPSC, AICS-0011, Allen Institute) were differentiated into cardiomyocytes (iPSC-CM) as recommended and then iPSC-CM were treated with endothelin (100nM, Tocris 1160) or endothelin+17β-estradiol (100 nM endothelin, 100 nM 17β-estradiol) overnight. iPSC-CMs were fixed with 4% paraformaldehyde, treated with 1% Triton X100, washed/blocked and incubated with primary antibodies to β-tubulin (Abcam:ab6046) or detyrosinated α-tubulin (Abcam:AB48389). Cells were washed, incubated with Alexa-568 secondary antibody, and imaged. Adult male Sprague Dawley rats were subjected to pulmonary artery banding (PAB) using an 18-gauge needle. Two weeks post banding, rats were randomly allocated to daily 17β-estradiol treatment (0.2 mg/kg PAB-E2) or ethanol vehicle (PAB-Vehicle) via intraperitoneal injection for two weeks. Formalin sections of right ventricular (RV) free walls were subjected to heat-mediated antigen retrieval (Reveal Decloaker, Biocare Medical), stained with primary antibodies to β-tubulin (Abcam:AB6046), desmin (ProSci:46-777), connexin-43 (Abcam:AB235585), and then counterstained with AlexaFluor-568 anti-rabbit secondary antibody (ThermoScientific:A-11036), Wheat Germ Agglutinin (WGA) AlexaFluor-488 (ThermoFisher:W21404), and mounted in ProLong Glass with NucBlue Antifade Mountant (ThermoFisherScientific:P36985). Confocal micrographs were collected on a Zeiss LSM900 Airyscan 2.0 microscope. Confocal images were blindly analyzed by RM using FIJI. Echocardiography and invasive closed-chest hemodynamics evaluated rodent right ventricular systolic pressure and RV function^5^. Animal studies were approved by the UMN Institutional Animal Care and Use Committee. Statistical analyses were performed using GraphPad Prism 10.1.

We first demonstrated 17β-estradiol directly impacted microtubule dynamics as 17β-estradiol significantly slowed microtubule polymerization rates *in vitro* (**Figure A**). Next, we showed the microtubule-suppressing effects of 17β-estradiol were present in human iPSC-CM to provide cardiac-cell relevance to our *in vitro* finding. As compared to control cells, endothelin treatment increased the density of both total and detyrosinated microtubules (**Figure B and C**). However, 17β-estradiol treatment blunted endothelin-mediated densification of total and detyrosinated microtubules (**Figure B and C**). Thus, 17β-estradiol reduced microtubule polymerization rates and counteracted chemical stress-induced microtubule remodeling in iPSC-CMs.

Then, we quantified the physiological effects of 17β-estradiol using both echocardiography and right heart catheterization using a translational approach by starting treatment two weeks after pulmonary artery banding to allow for the development of RV dysfunction (**Figure D**). Echocardiographic analysis revealed 17β-estradiol increased tricuspid plane annular systolic excursion (TAPSE) and right ventricular free wall thickening (**Figure E**). Importantly, right ventricular systolic pressure was equivalently elevated in PAB-Vehicle and PAB-E2 animals (**Figure E**). These data suggested 17β-estradiol imparted an inotropic effect independent of RV afterload.

**Figure 1.**
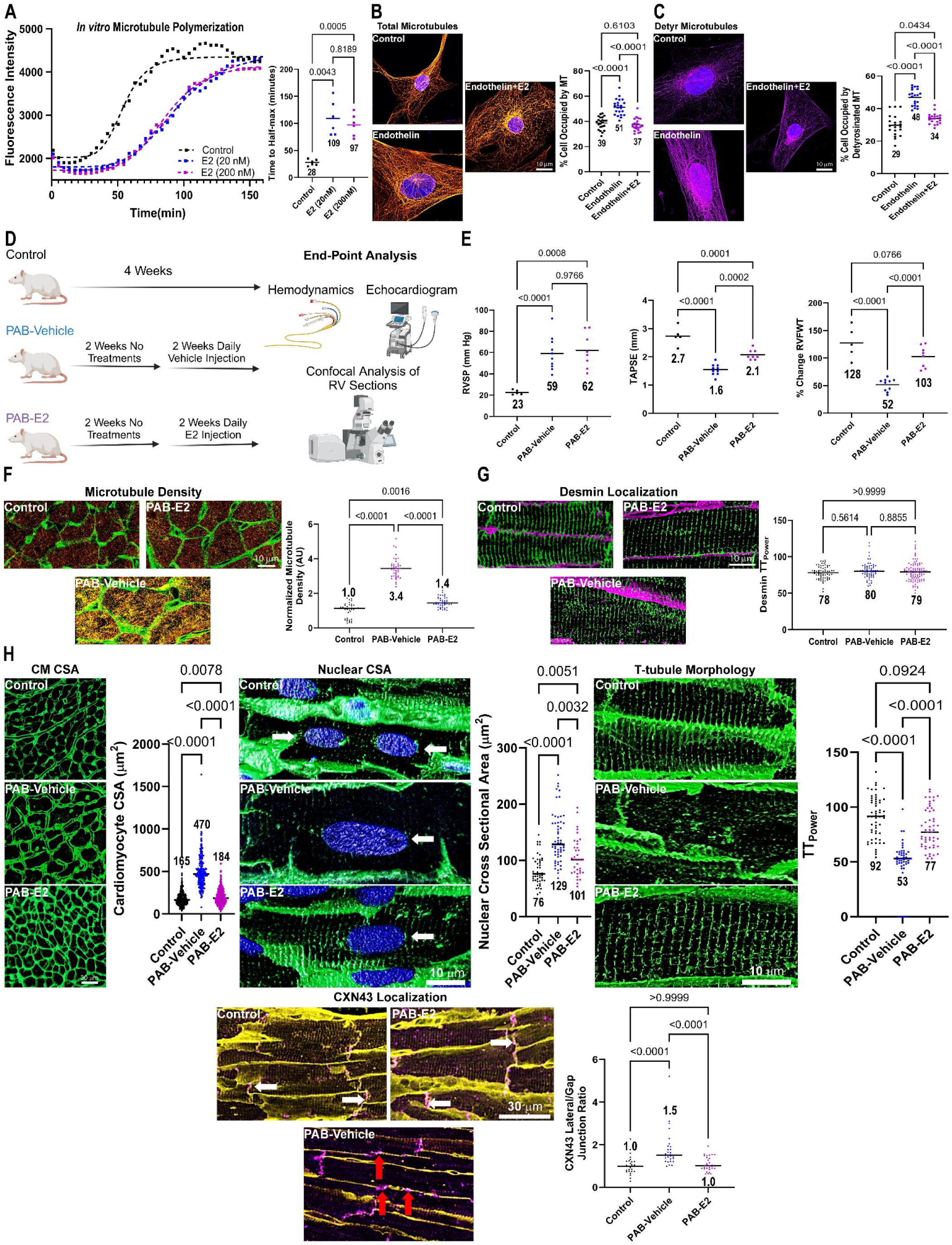
17β-Estradiol Limits Pathological Microtubule Stabilization and Improves Cardiac Function in Pulmonary Artery Banded Rats. (A) Microtubule polymerization kinetics in the presence of absence of 17β-estradiol. Quantification of time to half max in presence of vehicle, 20 nM 17β-estradiol, and 200 nM 17β-estradiol. *p*-values determined by one-way ANOVA with Tukey’s multiple comparisons test. Representative confocal micrographs of iPSC-CM stained with β-tubulin (orange, B), detyrosinated α-tubulin (purple, C), and DAPI (blue). Quantification of total microtubule (B) and detyrosinated microtubule (C) density. *p*-values determined by one-way ANOVA with Tukey’s multiple comparisons test for total microtubule density and Brown-Forsythe ANOVA and Dunnett’s T3 multiple comparisons test for detyrosinated microtubule density. (D) Representation of experimental approach for evaluating *in vivo* microtubule effects of 17β-estradiol. (E) E2 treatment did not reduce right ventricular systolic pressure but increased RV function determined by TAPSE and % change in RV free wall thickness. *p*-values determined by one-way ANOVA with Tukey’s multiple comparisons test for TAPSE and percent RV free wall thickening and Brown-Forsythe ANOVA and Dunnett’s T3 multiple comparisons test for RVSP. (F) WGA (green) and microtubule staining (orange) of RV free wall sections from control, PAB-Vehicle, and PAB-E2 RV sections with quantification of relative microtubule fluorescence intensity. *p*-values determined by one-way ANOVA with Tukey’s multiple comparisons test. (G) Confocal micrographs of RV sections stained with WGA (purple) and desmin (green). Desmin localization patterns were similar in all three experimental groups. *p*-values determined by Kruskal-Wallis and Dunn’s multiple comparisons test. (H) WGA (green) and DAPI (blue) delineated cardiomyocyte and nuclear size (white arrow). *p*-values determined by Kruskal-Wallis test with Dunn’s multiple comparison test. 17β-estradiol restored the normal striated t-tubule architecture in RV sections (*p*-values determined by Kruskal-Wallis test with Dunn’s multiple comparison test). 17β-estradiol prevented lateralization of connexin-43 (purple) in the RV (red arrows: lateralized connexin-43, white arrows: connexin-43 at intercalated disc). *p*-values determined by Kruskal-Wallis test with Dunn’s multiple comparison test.

Finally, we determined how 17β-estradiol treatment modulated RV cardiomyocyte microtubule phenotypes in PAB rats to assess if the observed improvements in cardiac function were associated with alterations in the microtubule cytoskeleton. Confocal microscopy revealed a significant increase in microtubule density in RV cardiomyocytes following PAB, which 17β-estradiol mitigated (**Figure F**). Importantly, 17β-estradiol did not alter desmin localization patterns (**Figure G)**, which implied its effects on cardiac morphology were predominately due to microtubule modulation rather than desmin regulation. The reduction in microtubule density with 17β-estradiol treatment limited both cardiomyocyte and cardiomyocyte nuclear hypertrophy (**Figure H**). 17β-estradiol also combatted other pathogenic microtubule-associated phenotypes as it improved t-tubule structural integrity, and prevented the lateral membrane localization of connexin-43 (**Figure H**). Thus, the cardioprotective effects of 17β-estradiol were paired with corrections of multiple microtubule-based phenotypes in RV cardiomyocytes.

In conclusion, we show 17β-estradiol slows microtubule polymerization *in vitro*, and counteracts endothelin-mediated microtubule densification in human iPSC-CM. In PAB rats, 17β-estradiol mitigates microtubule remodeling, and downstream pathogenic phenotypes in RV cardiomyocytes. Importantly, these structural changes occur without significant modulation of desmin localization, which further supports 17β-estradiol’s cellular effects are related to its microtubule-modulating properties. The correction of microtubule-associated cellular phenotypes ultimately manifest as improvements in RV function independent of afterload. In summary, our data further support a pathogenic role of hyperstabilized microtubules in cardiac dysfunction. We therefore nominate the direct modulation of microtubule remodeling as a novel mechanism by which 17β-estradiol exerts cardioprotective effects in addition to its known signaling roles^4^.

